# A role for DIS3L2 over human nonsense-mediated mRNA decay targets

**DOI:** 10.1101/722702

**Authors:** Paulo J. da Costa, Juliane Menezes, Margarida Saramago, Juan F. García-Moreno, Hugo A. Santos, Margarida Gama-Carvalho, Cecília M. Arraiano, Sandra C. Viegas, Luísa Romão

**Affiliations:** Department of Human Genetics, Instituto Nacional de Saúde Doutor Ricardo Jorge, Lisboa, Portugal; University of Lisboa, Faculty of Sciences, BioISI - Biosystems and Integrative Sciences Institute, Lisboa, Portugal; Instituto de Tecnologia Química e Biológica António Xavier, Universidade Nova de Lisboa, Oeiras, Portugal

## Abstract

The nonsense-mediated decay (NMD) pathway selectively degrades mRNAs carrying a premature translation-termination codon but also regulates the abundance of a large number of physiological mRNAs that encode full-length proteins. In human cells, NMD-targeted mRNAs are degraded by endonucleolytic cleavage and exonucleolytic degradation from both 5’ and 3’ ends. This is done by a process not yet completely understood that recruits decapping and 5’-to-3’ exonuclease activities, as well as deadenylating and 3’-to-5’ exonuclease exosome activities. In yeast, DIS3/Rrp44 protein is the catalytic subunit of the exosome, but in humans, there are three known paralogues of this enzyme: DIS3, DIS3L1, and DIS3L2. DIS3L1 and DIS3L2 exoribonucleases localize in the same compartment where NMD occurs, but little is known about their role in this process. In order to unveil the role of DIS3L2 in NMD, here we show that some NMD-targets accumulate in DIS3L2-depleted cells. mRNA half-life analysis further supports that these NMD-targets are in fact DIS3L2 substrates. Besides, we observed that DIS3L2 acts over full-length transcripts, through a process that also involves UPF1. Moreover, DIS3L2-mediated decay is dependent on the activity of the terminal uridylyl transferases Zcchc6/11 (TUT7/4). Together, our findings establish a role for DIS3L2 and uridylation in NMD.

## INTRODUCTION

In eukaryotes, mRNAs are subjected to quality control mechanisms [1–4]. Nonsense-mediated decay (NMD) is an mRNA surveillance mechanism that recognizes and rapidly degrades eukaryotic mRNAs harboring premature translation-termination codons (PTCs) [2,3]. However, it has been shown that NMD also targets normal (wild-type) transcripts that have “NMD-inducing features”, including transcripts containing upstream open reading frames (uORFs) in the 5’ untranslated region (UTR), long 3’ untranslated regions (3’UTRs), introns in the 3’UTR located more than 50 nucleotides (nts) downstream from the stop codon, selenocysteine codons, or products of alternative splicing that contain PTCs [5–9]. In fact, NMD controls the steady-state levels of about 10% of the eukaryotic mRNAs [10–13].

NMD is dependent on the UPF1 protein, being triggered by its phosphorylation [14–16]. The UPF1 phosphorylation is mediated by the SMG1 kinase upon interaction between the UPF1, the translation-termination factors eRF1 and eRF3, and UPF2 present in the exon junction complexes of the mRNA [14–16]. This event promotes the mRNA degradation that involves both exonucleolytic and endonucleolytic pathways in mammalian cells [17]. The different decay paths all seem to require phosphorylated UPF1, which interacts with SMG6, SMG5-SMG7 and Proline-Rich Nuclear Receptor Coactivator 2 (PNRC2) [18–21]. The first pathway relies on the recruitment of SMG5-SMG7 heterodimer or SMG5-PNCR2 that recruit the decapping enzymes (DCP1 and DCP2) and the CCR4-NOT deadenylation complex. Hence, the mRNA is deadenylated and decapped allowing the 5’-to-3’ and 3’-to-5’ mRNA degradation by the exoribonuclease 1 (XRN1) and the RNA exosome, respectively [18,19,22–27]. Of note, Nicholson and colleagues (2018) have found no evidence for the existence of a physical or functional interaction between SMG5 and PNRC2. Instead, authors have shown that UPF1 interacts with PNRC2 and that it triggers 5’-to-3’ exonucleolytic decay [21]. On the other hand, the endonucleolytic-dependent pathway starts with an endonucleolytic cleavage mediated by the SMG6 protein [28,29] that generates two mRNA intermediates with unprotected ends that are then degraded [30,31]. Despite the differences between the NMD branches, several studies have pointed out a high degree of redundancy of the affected transcripts in both pathways [7,8,20,32].

In humans, the mRNA degradation in the cytoplasm is achieved either 5’-to-3’ by the XRN1 or 3’-to-5’ by the DIS3 protein family: DIS3, DIS3L1 and DIS3L2. DIS3 and DIS3L1, as members of the RNA exosome, were implicated in normal mRNA decay, as well as in NMD and non-stop decay (NSD) [33–36]. DIS3L2, unlike its family counterparts, is not part of the exosome and has been associated with an uridylation-dependent RNA degradation pathway [37–43]. This pathway relies on the addition of untemplated uridine residues to mRNAs, tRNAs, miRNAs, snRNAs, among other classes of RNAs, by proteins called terminal uridylyl transferases (TUTases or TUTs) [40,42].

Here, we report the involvement of DIS3L2 in the degradation of natural NMD targets. Our results show that some NMD-targets accumulate in DIS3L2-depleted cells. In addition, mRNA half-life analyses indicate that these NMD-targets are in fact substrates for DIS3L2. We also observed that DIS3L2 acts over full-length transcripts, along with UPF1, being DIS3L2-mediated decay dependent on the activity of the terminal uridylyl transferases Zcchc6/11 (TUT7/4).

## MATERIALS AND METHODS

### Plasmid constructs

The pTRE2pur vector (BD Biosciences) containing each of the human β-globin gene variants βWT (wild type), β15 (PTC at codon 15; TGG→TGA), β26 (PTC at codon 26; GAG→TAG), β39 (PTC at codon 39; CAG→TAG) have been used before in our laboratory and were obtained as previously described [44–46].

### Cell culture, plasmid and siRNA transfections

HeLa and HEK293 cells were grown in Dulbecco’s modified Eagle’s medium (DMEM) supplemented with 10% fetal bovine serum. Transient transfections of siRNAs were carried out using Lipofectamine 2000 reagent (Invitrogen) according to the manufacturer’s instructions in 35-mm plates using 200 or 30 pmol of siRNA oligonucleotides and 4 μl of transfection reagent. Twenty-four hours after first siRNA transfection, cells were transfected with additional 50 pmol of siRNAs and 500 ng of the tested construct DNA (when necessary). The siRNA oligonucleotides used for transfections were designed with 3’-dTdT overhangs, and purchased as annealed, ready-to-use duplexes from Thermo Fisher Scientific. All sequences are available in Table S1. Twenty-four hours later, cells were harvested for RNA and protein expression analysis. To perform the mRNA half-life analyses, after this additional 24 hours, HeLa cells were treated with 60 μM of the adenosine analogue 5,6-dichloro-1-3-D-ribofuranosylbenzimidazole (DRB; Sigma-Aldrich®) to inhibit transcription and RNA was extracted at different time points for further RT-qPCR analysis.

### Isolation of total RNA and protein lysates

Total RNA and protein extracts were obtained from lysis of transfected cells. Briefly, cells were lysed with 100 μl of NP40 buffer [50 mM Tris-HCl pH=7.5, 10 mM MgCl_2_, 100 mM NaCl, 10% (v/v) glycerol and 1% (v/v) Nonidet P-40 (Roche)], supplemented with Proteinase and RNAse inhibitors. After lysis, samples were spun down at maximum speed for 1 minute. Twenty microliters have been collected for protein analysis and RNA was extracted from remaining supernatant with the Nucleospin RNA extraction II Kit (Macherey-Nagel).

### Western blot analysis

Protein lysates mixed with 1x SDS-PAGE Sample Loading Buffer (NZYtech) were denatured at 95°C for 10 minutes and centrifuged at 500g for 5 minutes. Then, they were resolved in a 10% SDS-PAGE gel and transferred to polyvinylidene difluoride (PVDF) membranes (Bio-Rad), according to standard protocols. After transfer, nonspecific sites were blocked with blocking buffer [TBS-0.5% TW20: 50 mM Tris-HCl pH7.5, 150 mM NaCl, 5% non-fat dried milk, and 0.5% (v/v) Tween 20] for 1 hour at room temperature (RT). The membrane was incubated overnight (O/N) at 4°C with mouse anti-α-tubulin (Roche, loading control) at 1:4000, rabbit anti-DIS3L2 (Novus Biologicals) at 1:200, rabbit anti-XRN1 (Novus Biologicals) at 1:500 or mouse anti-FLAG M2 (Sigma) at 1:1000 dilution in blocking buffer. After O/N incubation, the membrane was rinsed 4×5 minutes, with large volumes of wash buffer [TBS-0.05% TW20: 50 mM Tris-HCL pH7.5, 150 mM NaCl and 0.05% (v/v) Tween 20]. Afterwards, detection was carried out by incubating the membrane with the secondary antibody goat anti-rabbit horseradish peroxidase conjugate (Sigma) diluted 1:3000 (or 1:4000 goat anti-mouse (Bio-Rad) for α-tubulin) in 12 ml of blocking buffer for 1 hour at RT and followed by enhanced chemiluminescence.

### Reverse transcription-coupled quantitative PCR (RT-qPCR)

First-strand cDNA was synthesized from 1 μg of total RNA using reverse trancriptase (NZYtech) according to the manufacturer’s instructions. Real time PCR was performed with the ABI7000 Sequence Detection System (Applied Biosystems) using SYBR Green PCR Master Mix (Applied Biosystems). The relative expression levels of β-globin mRNA were normalized to the internal control GAPDH mRNA in HeLa cells and calculated using the comparative Ct method (2^−ΔΔCt^) [47]. The Ct values of variant β-globin mRNA amplicons were compared to the respective βWT counterpart or to βWT at LUC siRNA conditions, as indicated in Figures and normalized with the reference amplicon Ct value. The amplification efficiencies of each primer pair were determined by dilution series. The forward and reverse primer sequences are available in Table S2. Technical triplicates from three independent experiments were assessed in all cases. To check for DNA contamination, quantitative PCR without reverse transcription was also performed.

### Northern blot analysis

For Northern blot analysis, equal amounts of human total RNA samples (15 µg) were separated under denaturing conditions in an agarose MOPS/formaldehyde gel (1%) following the procedure described in [38]. Briefly, the RNA was transferred onto Hybond-N+ membranes by capillarity using 20×SSC as transfer buffer, and UV cross-linked to the membrane immediately after. Membranes were then hybridized, in PerfectHyb Buffer (Sigma) for 16 hours, with the specific riboprobe at 68°C or oligoprobe at 43°C. After hybridization, membranes were rinsed at RT in a 2X SSC/0.1% SDS solution, followed by washing in three subsequent 15 minutes steps in SSC (2X, 1X or 0.5X, respectively)/0.1% SDS solutions at the hybridization temperature. Signals were visualized by Phosphor Imaging (FUJI TLA-5100 Series, Fuji) and analyzed using the ImageQuant software (GE Healthcare). Riboprobe templates were generated by standard PCR with specific primers and carried out on human cDNA. The *in vitro* transcription reaction was performed in the presence of an excess of [32P]-α-UTP over unlabeled UTP, using the T7 polymerase from Promega (a T7 RNA polymerase promoter sequence was added in the template DNA by the antisense primer). 28S rRNA was detected using an oligoprobe. The riboprobe was purified with a G50 column (GE Healthcare) and the oligoprobe with a G25 column (GE Healthcare) to remove unincorporated nucleotides prior to hybridization (see primers used for probes design on Supporting Table S3).

### RNA 3’-RACE

We used optimized 3’-RACE method (PASE method) according to the protocol described in ref. [48]. Total RNA (500 ng) from cells treated with DIS3L2/XRN1 and LUC siRNAs was ligated to an adaptor (Linker Oligo) at 4°C for 12 hours with 40 units of T4 RNA ligase (Fermentas). RNA was used as template for RT-PCR with an adapter-specific primer (BRevOligo) and 10 units of Transcriptor Reverse Transcriptase (Roche) according to the manufacturer’s instructions. The products of reverse transcription were amplified using 2 μl aliquot of the RT-PCR reaction, 20 pmol of a gene specific primer (GADD45A External) and an adapter-specific BRevOligo primer, 250 μM of each dNTP, 1.25 units of DreamTaq (Fermentas) and 1x DreamTaq buffer. Cycling conditions were as follows: a step at 95°C/10 min; 30 cycles of 95°C/1 min, 45°C/1.5 min, 72°C/3 min; and a final 72°C/10 min step. PCR reactions were 1:20 diluted in H_2_O, and 2 μl aliquot were used for Nested PCR with 20 pmol of an internal primer (GADD45A Internal) and an adapter-specific ARevOligo primer. The same PCR and cycling conditions were maintained. Products were cloned into pGEM-T Easy vector (Promega). Bacterial colonies obtained after transformation were screened by colony PCR for the presence of inserts of appropriate size. The plasmids with inserts of interest were extracted (NZYMiniprep, Nzytech) and sequenced using M13 FW and M13 REV universal primers. Primers used are listed in Table S3.

### Bioinformatic analysis

Transcriptome data for DIS3L2 KD in HeLa cells was retrieved from the supplementary material provided by Lubas *et al*. (2014) and refers to a list of 873 genes identified by differential expression (DE) analysis using the DESeq algorithm [[34]; Table S4)]. A dataset of UPF1 KD in HeLa cells was produced by Tani *et al*. [49]. To generate a list of genes that can be directly compared to the DIS3L2-dependent transcriptome, raw data for Illumina paired-end mRNA-seq were retrieved from SRA (identifiers DRS001615 - DRS001618) and analyzed as described by Lubas *et al*. [34]. Results obtained are presented in Table S4.

### Statistical analysis

When appropriate, Student’s unpaired, two-tailed t-test was used for estimation of statistical significance. Significance for statistical analysis was defined as p<0.05 (*), p<0.01 (**) and p<0.001(***). Results are expressed as mean ± standard deviation from at least three independent experiments.

## RESULTS

### DIS3L2 knockdown does not affect levels of nonsense-mutated human β-globin transcripts

In order to unveil the role of DIS3L2 in NMD, we knocked-down (KD) this ribonuclease in HeLa cells and transiently transfected them with constructs containing different human β-globin variants: βWT, β15, β26, β39 [44]. After monitoring the KD efficiency by Western blot (Fig 1A), the impact of altering DIS3L2 levels on the expression of the various reporter β-globin mRNAs was monitored by Northern blot (Fig S1) and quantitative RT-qPCR (Fig 1B). As a control, cells were treated with Luciferase (LUC) siRNA. In control conditions, without DIS3L2 KD, the mRNA levels of β15 were close (1.22-fold) to the βWT mRNA levels (arbitrarily set to 1), and both NMD-sensitive variants were significantly reduced, as previously shown [44]. In conditions of DIS3L2 KD, none of the β-globin variants mRNA levels changed significantly: the βWT mRNA levels had a 1.34-fold increase, β15 was 1.43-fold higher, β26 increased 1.19-fold, and β39 1.13-fold, when compared to βWT in control conditions (Fig 1B). Thus, these results indicate that DIS3L2 exoribonuclease does not seem to be involved in the general mRNA turnover or NMD of human β-globin transcripts.

**Fig 1.**
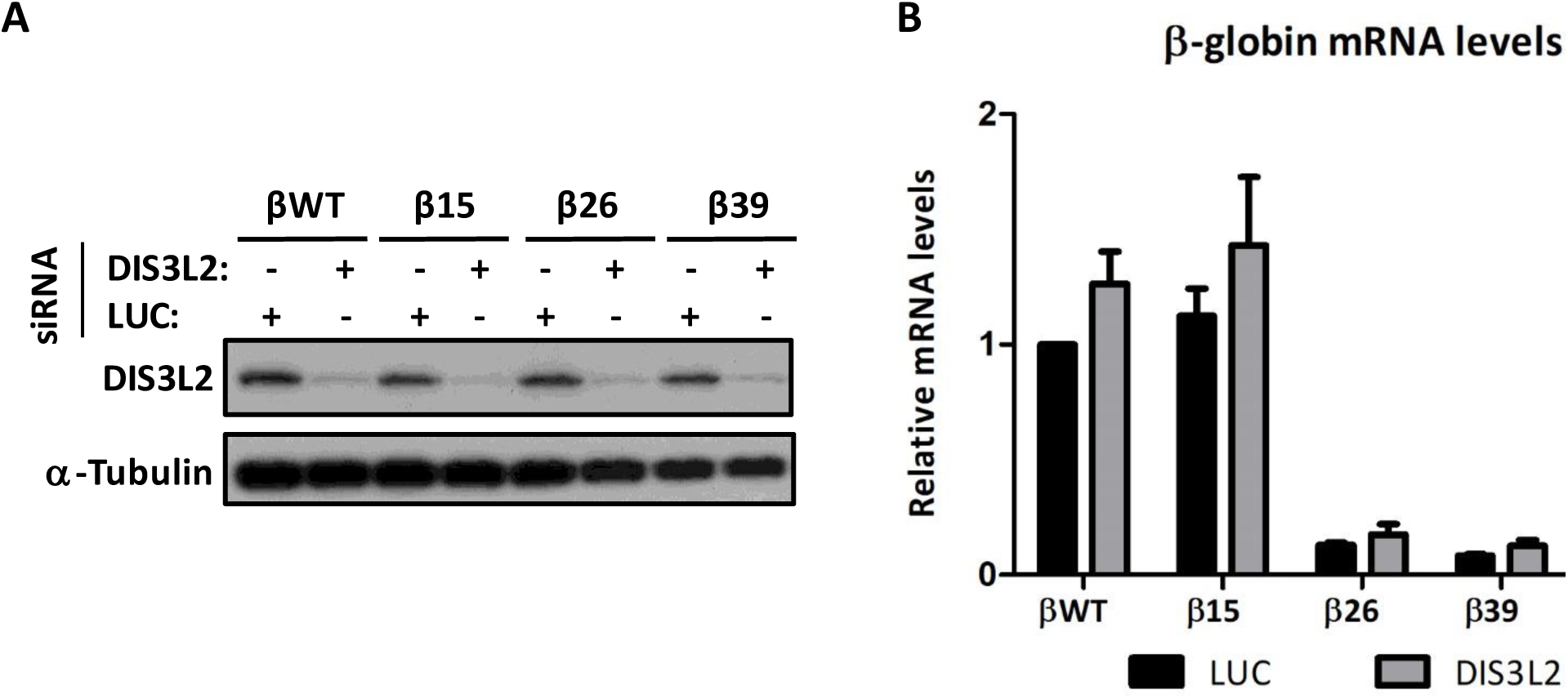
DIS3L2 depletion does not affect levels of the normal or nonsense-mutated human β-globin transcripts expressed in HeLa cells. (A) Representative Western blot analysis of HeLa cells extracts transfected (+) with control Luciferase siRNA (LUC siRNA), or with siRNA targeting the human DIS3L2. After siRNA treatment, cells were transfected with the plasmids expressing wild type (βWT), NMD-resistant (β15), or NMD-sensitive (β26, β39) human β-globin mRNAs. Protein levels present in the cell extracts were analyzed by Western blot for DIS3L2 and α-tubulin (loading control), to monitor endogenous DIS3L2 knockdown. Identification of each band is indicated to the left of the gel image. (B) Histogram represents fold-change of each sample relative to the control [βWT at Luciferase (LUC) siRNA-treated cells], arbitrarily set to 1. mRNA levels were determined by RT-qPCR using primers specific for human β-globin gene, and for glyceraldehyde-3-phosphate dehydrogenase gene, and compared to the βWT mRNA levels (defined as 1). Average and standard deviation (SD) of three independent experiments corresponding to three independent transfections are shown in the histogram.

### UPF1 and DIS3L2 can act in common transcripts

Despite the fact that our results do not suggest a connection between DIS3L2 and normal or nonsense-mutated β-globin mRNA decay, we raised the hypothesis that this ribonuclease could function in a transcript specific manner. Considering this, we selected 12 well characterized natural NMD-targets from the literature [SMG5 [11,13,50,51], ARFRP1 [11], BAG1 [50], ANTXR1 [11], GABARAPL1 [51], SMG1 [11,13,50], SLC7A11 [52], SLC1A3 [11], GADD45A [11,54], GADD45B [10], PLXNA1 [11], and ATF3 [10]] and proceeded to the analysis of their mRNA levels, with and without DIS3L2 depletion in HeLa cells. As a positive control for this experiment, we knocked-down UPF1. After monitoring the KD efficiency by Western blot (Fig 2A), the impact of altering DIS3L2 levels on the expression of the 12 selected natural NMD targets was assessed by RT-qPCR (Fig 2B-H and S2B-F). As expected for NMD-targets, we observed a significant increase in the levels of all studied transcripts, upon UPF1 depletion, compared to the levels observed in control conditions [luciferase (LUC) siRNA-treated cells], arbitrarily set to 1. DIS3L2 KD does not affect levels of the SMG5 (Fig 2B), GABARAPL1 (Fig 2C), ARFRP1 (Fig S2B), BAG1 (Fig S2C), and ANTXR1 (Fig S2D) mRNAs. The other natural NMD-targets (SMG1, SLC7A11, SLC1A3, GADDD45A, ATF3, GADD45B, and PLXNA1) significantly accumulate upon DIS3L2 depletion (Fig 2E-H and Fig S2E-F). Together, these results indicate that DIS3L2 might be involved in NMD. Thus, considering this interesting and unexpected result, we proceeded to investigate how the DIS3L2 functions in the NMD mechanism.

**Fig 2.**
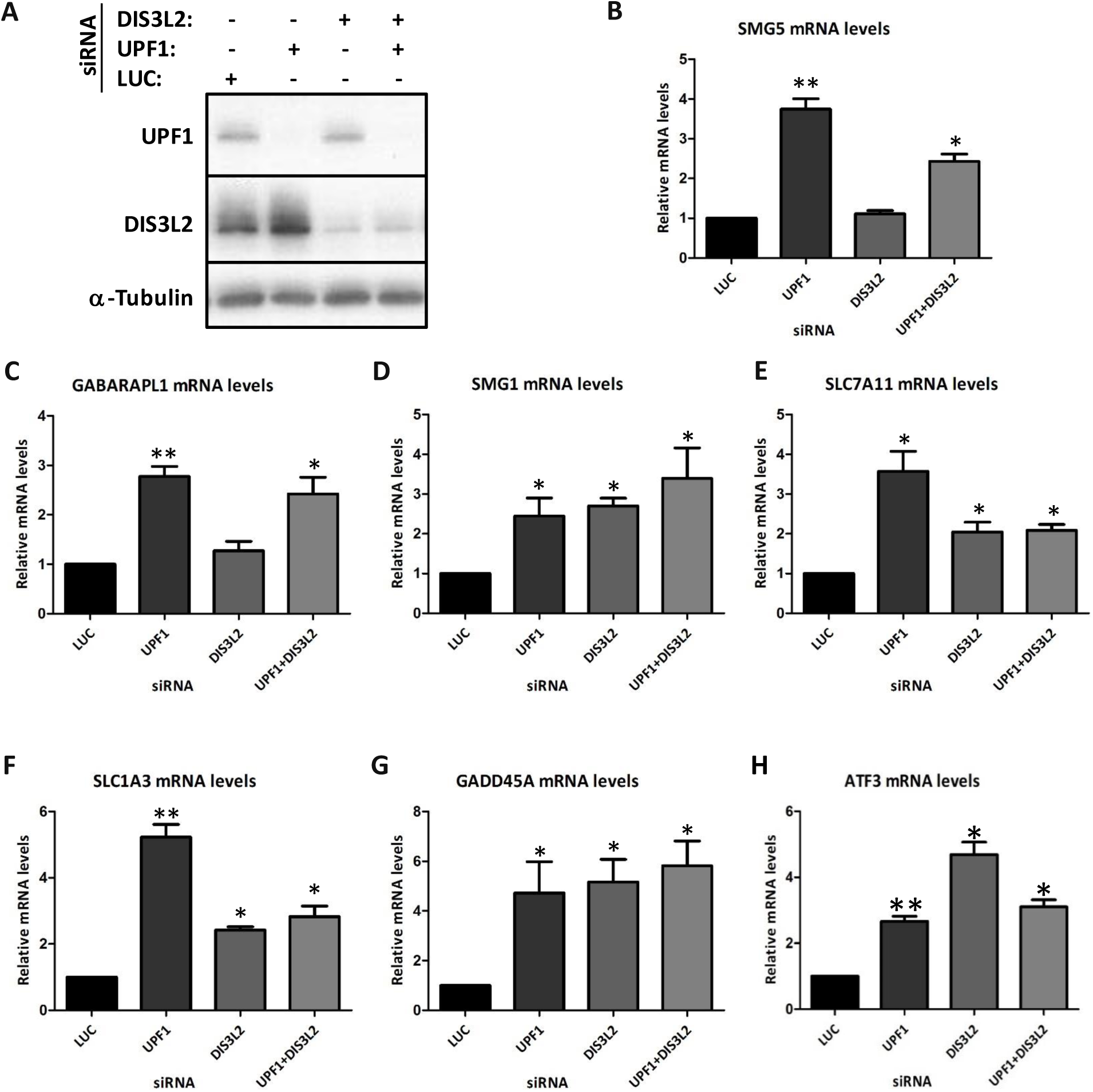
UPF1 and DIS3L2 seem to function in the same pathway. (A) Representative Western blot analysis of HeLa cells extracts transfected with (+) control Luciferase (LUC), UPF1, DIS3L2 or DIS3L2 plus UPF1 siRNAs. Protein levels present in the cell extracts were analyzed by Western blot for UPF1, DIS3L2, and α-tubulin (loading control) to monitor UPF1 and DIS3L2 knockdown. Identification of each band is indicated to the left of the gel image. (B-H) mRNA levels of natural NMD-targets (SMG5, GABARAPL1, SMG1, SLC7A11, SLC1A3, GADD45A, and ATF3) were determined by RT-qPCR using primers specific for each case (as indicated above each histogram), and for glyceraldehyde-3-phosphate dehydrogenase (GAPDH) gene. Histograms represent fold-change of each studied NMD-target in UPF1- or DIS3L2-depleted HeLa cells, relative to its level at Luciferase (LUC) siRNA-treated cells, arbitrarily set to 1. Average and standard deviation (SD) of three independent experiments corresponding to three independent transfections are shown in the histograms. Asterisks (*) indicate statistical significance relative to the control conditions for each NMD-target. *P<0.05, **P<0.01. Asterisks (*) indicate statistical significance relative to the mRNA levels in control conditions (LUC siRNA-treated cells). \**P*<0.05, \*\**P*<0.01.

### UPF1 and DIS3L2 function in the same pathway

To investigate whether DIS3L2 and UPF1 function in the same pathway, we accessed the mRNA levels of some of the endogenous NMD targets (SMG5, GABARAPL1, SMG1, SLC7A11, SLC1A3, GADD45A, and ATF3) studied in this work in a double knockdown of UPF1 and DIS3L2. Results were compared to those obtained in single DIS3L2 or UPF1 KD (Fig 2B-H). In the case of SMG5 mRNA, and comparing to the control conditions, we observed a significant 3.7-fold increase in cells with UPF1 KD, as expected for an NMD-target. Despite the DIS3L2 KD does not affect its levels, the double DIS3L2+UPF1 KD induces a significant 2.4-fold increase in the SMG5 mRNA levels; however, this increase is lower than that observed upon UPF1 KD (Fig 2B). Regarding the GABARAPL1 mRNA (Fig 2C), mRNA levels do not significantly change upon DIS3L2 KD. In cells with double KD, we observed a significant 2.4-fold increase in the GABARAPL1 mRNA levels, similar to the significant 2.8-fold increase observed upon UPF1 KD. Taking these results together, GABARAPL1 mRNA is an NMD-target, but, DIS3L2 is not significantly involved in its degradation. Regarding the SMG1, SLC7A11, SLC1A3, GADD45A, and ATF3 mRNAs (Fig 2D-H), we observed a significant increase in UPF1-silenced cells. Moreover, the SMG1 mRNA levels significantly increased 2.7-fold and 3.4-fold, in cells with a single DIS3L2 KD or a double KD of DIS3L2 and UPF1, respectively (Fig 2D). Despite being higher than that one with single knockdown, SMG1 mRNA levels upon double knockdown do not show a significant additive effect. Indeed, this difference is far from being statistically significant (p-value>0.5) (Fig 2D). Concerning the SLC7A11 mRNA (Fig 2E), upon DIS3L2 KD, we observed a significant 2.0-fold increase, and in cells with the double KD, we observed a significant 2.1-fold increase (Fig 2E). Therefore, for SLC7A11 mRNA is very clear that DIS3L2+UPF1 double knockdown has no additive effect. Considering the SLC1A3 mRNA (Fig 2F), we observed a significant 2.4-fold increase in cells with the DIS3L2 KD, and a similar significant 2.8-fold increase in cells with the double depletion. Thus, the DIS3L2+UPF1 double KD does not show an additive effect over SLC1A3 mRNA levels. Considering the GADD45A mRNA (Fig 2G), we observed a significant 5.2-fold increase upon DIS3L2 KD, and, in cells with the double KD, we observed a significant 5.8-fold increase in its mRNA levels. Once again, concerning the GADD45A mRNA levels, the DIS3L2+UPF1 double KD does not induce an additive effect when compared to the effect of the single DIS3L2 or UPF1 depletions. Concerning the ATF3 mRNA (Fig 2H), we observed a significant 4.7-fold increase in cells with the DIS3L2 KD, and upon the double knockdown, we observed a significant 3.1-fold increase, showing no additive effect of the double KD over ATF3 mRNA levels. Curiously, upon double UPF1+DIS3L2 KD, the mRNA levels of some NMD-targets (for example, SLC7A11 and SLC1A3) are similar to those obtained in the single KD with the weakest effect, indicating that UPF1- and DIS3L2-targets have different relative levels of sensitiveness to these proteins. Taken, together, these results indicate that DIS3L2 is involved in the degradation of NMD-targets through a process that also involves the UPF1 protein.

### DIS3L2 is involved in the decay of some NMD-targets

Next, to collect further evidence that the transcripts that respond to the DIS3L2 depletion are DIS3L2 targets, we analyzed the mRNA half-life of some of the endogenous NMD-targets (SMG1, SLC7A11, GADD45A and ATF3 mRNAs). This was done in control conditions, or in cells depleted of UPF1 or DIS3L2. For this, cells were treated with siRNAs targeting LUC (Control), UPF1 or DIS3L2, as before. Then, transcription was inhibited by the addition of the adenosine analogue 5,6-dichloro-1-3-D-ribofuranosylbenzimidazole (DRB) and the mRNA levels of the endogenous NMD-targets were measured before and at several time points after DRB addition. The KD efficiency in each time point was accessed by Western blot (Fig 3A).

**Fig 3.**
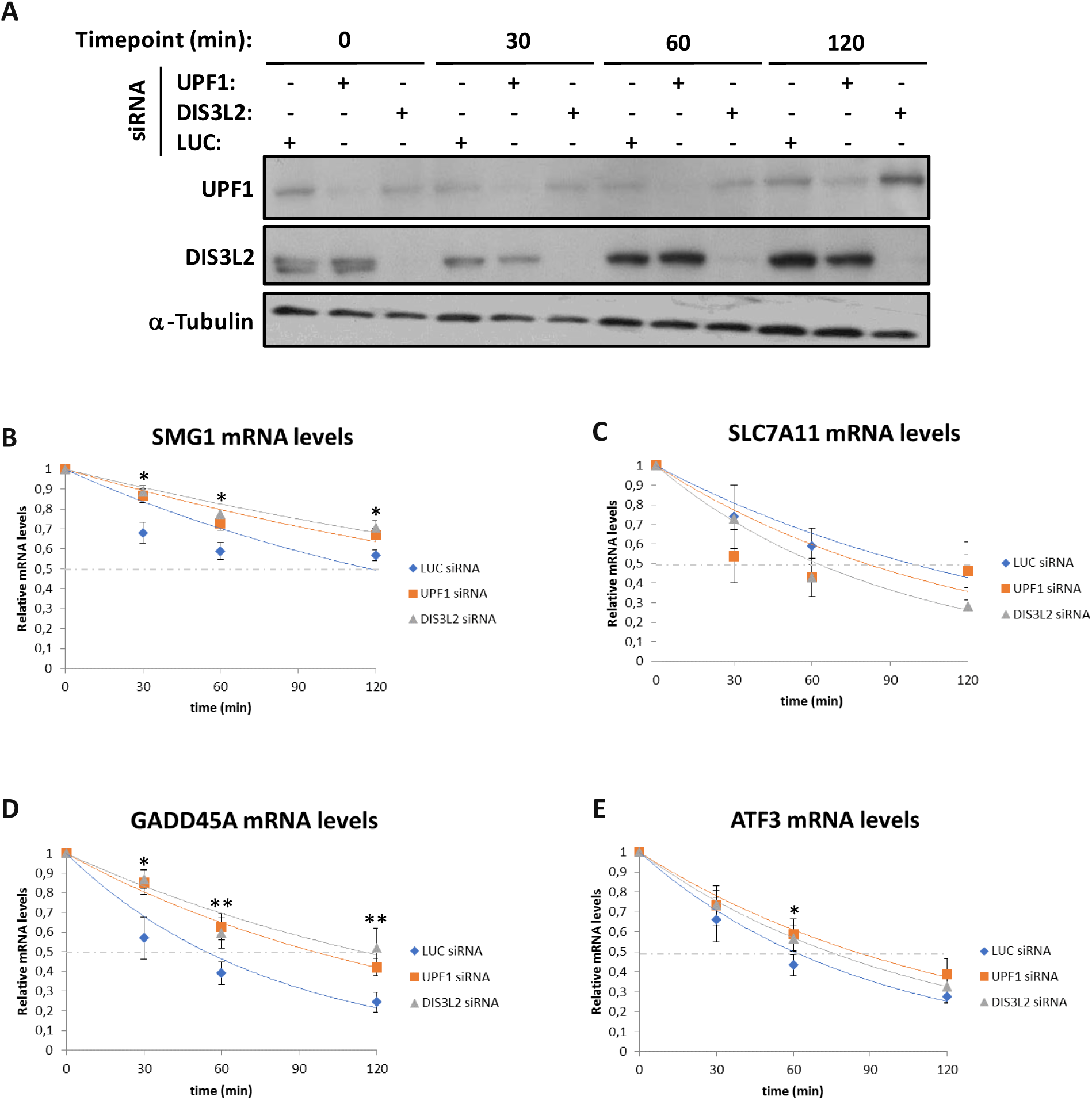
DIS3L2 is involved in the decay of some NMD-targets. (A) Representative Western blot analysis of HeLa cells extracts transfected with (+) or without (-) control Luciferase (LUC) siRNAs, or with siRNAs targeting the human UPF1 or DIS3L2. Protein levels present in the cell extracts were analyzed by Western blot for UPF1, DIS3L2 and α-tubulin (loading control) to monitor knockdown of UPF1 and DIS3L2, respectively. Identification of each band is indicated to the left of the gel image. (B-E) Cells were harvested, at various time points (0, 30, 60, 120 minutes) after treatment with the transcription inhibitor 5,6-dichloro-1-3-D-ribofuranosylbenzimidazole and mRNA quantified by RT-qPCR. To calculate the mRNA half-lives, each time point data was expressed as a ratio between the level of the NMD-target and glyceraldehyde-3-phosphate dehydrogenase (GAPDH) mRNA and normalized to the average value of all time points from a single transfection. The ratios were then renormalized to the average initial time point from all transfections (time 0 = 1). Each point represents the mean (n=3, at least). Standard deviations are also shown. Logarithmic regression analysis was performed to calculate the mRNAs half-lives. *P<0.05, **P<0.01.

Considering the SMG1 mRNA (Fig 3B), we observed that in control conditions (LUC siRNA-treated cells), its half-life is 118 minutes (min). Upon UPF1 or DIS3L2 KD, we observed a significant increase in the SMG1 mRNA half-live, as shown in Fig 3B (estimated to be far more than 120 min). Regarding the SLC7A11 mRNA (Fig 3C), we observed that in control conditions its half-life is 98 min. However, both UPF1 and DIS3L2 KDs decrease the SLC7A11 mRNA stability, being its half-live 80 and 62 min, respectively. This result indicates that SLC7A11 mRNA is not a direct target of UPF1 and DIS3L2. This was unexpected as our results show an increase in SLC7A11 mRNA levels in cells depleted of UPF1, DIS3L2, or UPF1 plus DIS3L2 (Fig 2E). Thus, we can speculate that SLC7A11 mRNA accumulation observed previously is due to an indirect effect. Concerning the GADD45A mRNA (Fig 3D), we observed that in control conditions (LUC siRNA-treated cells) its half-life is 54 min, while upon UPF1 or DIS3L2 KD we observed a significant increase in its half-live of 96 and 115 min, respectively. This result shows that GADD45A mRNA is also a target of UPF1 and DIS3L2. Regarding the ATF3 mRNA (Fig 3E), we observed that its half-life in control condition is 60 min, and upon UPF1 or DIS3L2 KD, it increases to 85 and 74 min, respectively. Despite not being significant, this difference indicates that the ATF3 mRNA is a target of both UPF1 and DIS3L2 factors. From these results we can conclude that DIS3L2 is involved in the decay of specific NMD substrates.

### Direct DIS3L2-mediated degradation of an NMD-target

To better understand the mechanism through which DIS3L2 is involved in NMD and considering that SMG6 is an endonuclease involved in NMD [28,29], we next tested whether DIS3L2 degrades intermediates that are first cleaved by this endonuclease. Bearing this in mind, we specifically analyzed by Northern blot (Fig 4) the GADD45A mRNA expressed in HeLa cells under control conditions (treated with LUC siRNAs) or treated with DIS3L2 siRNAs. GADD45A mRNA was analyzed using two different probes: one specific for the 5’UTR and another one for the 3’UTR. Using these probes, we observed that GADD45A mRNA accumulates in the full-length form and no decay intermediates were detected in the conditions tested. Furthermore, comparable GADD45A mRNA levels were observed with both probes. These results indicate that the GADD45A mRNA does not seem to be targeted by SMG6 endonuclease prior to the DIS3L2 degradation. To better confirm the absence of NMD-specific endonucleolytic cleavage sites, we further analyzed by Northern blot the GADD45A mRNA expressed in HeLa cells under control conditions (treated with LUC siRNAs), or treated with DIS3L2, XRN1, or DIS3L2+XRN1 siRNAs (Fig S3). In this case, GADD45A mRNA was analyzed with one probe specific for the GADD45A 3’UTR. Results show that GADD45A mRNA accumulates as full-length transcript and no significant 3’-intermediates are observed (Fig S3). Based on these results, it seems that the GADD45A mRNA is degraded without significant involvement of the SMG6 endonuclease prior to the DIS3L2 degradation. Together our results indicate that DIS3L2 might have a direct role on the degradation of NMD-targets.

**Fig 4.**
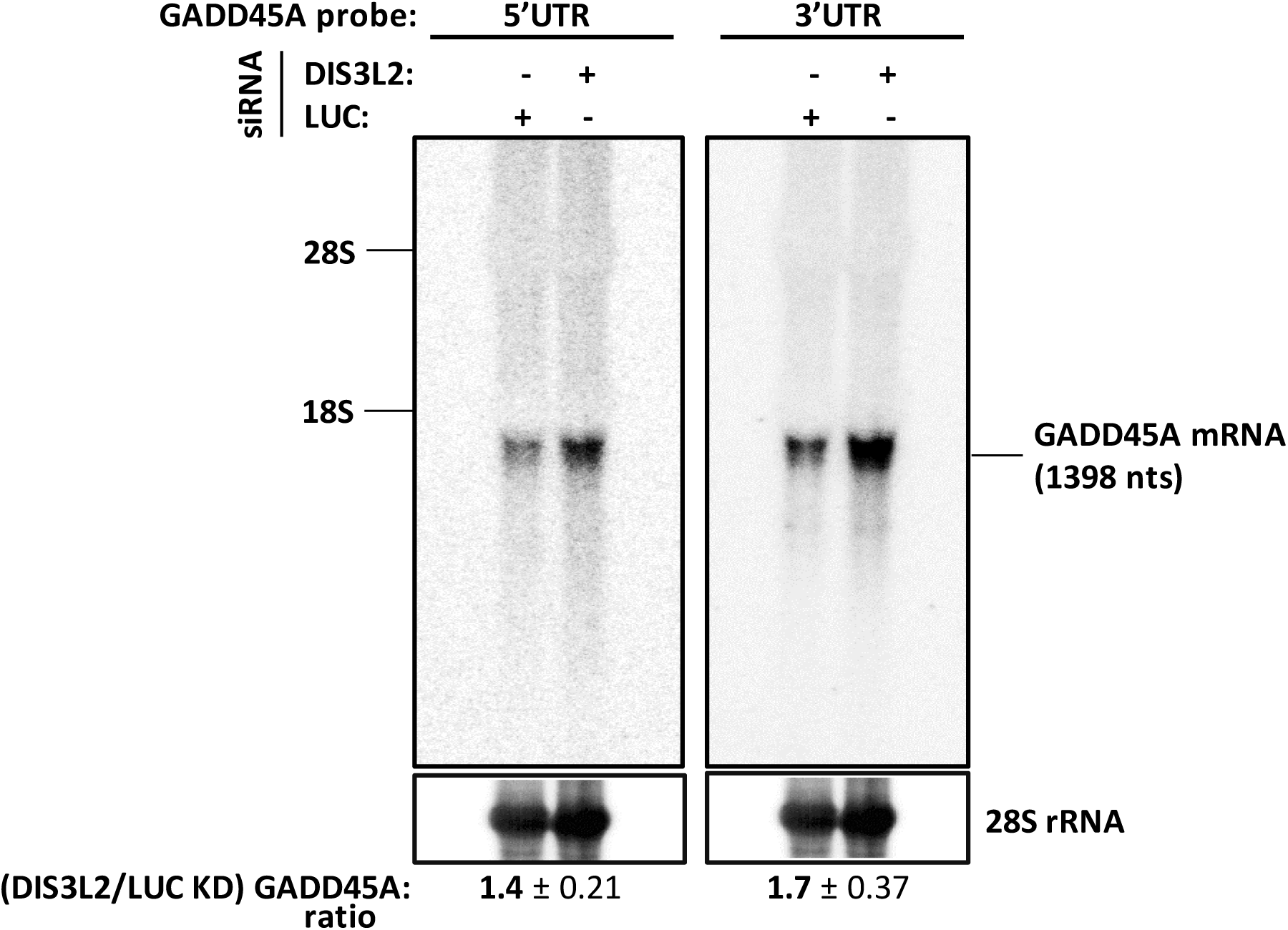
DIS3L2 targets full-length transcripts. Northern blot analysis of total RNA obtained from HeLa cells treated with (+) or without (-) siRNAs targeting DIS3L2. Fifteen micrograms of total RNA were separated under denaturing conditions on an agarose/MOPS/formaldehyde gel (1%) and GADD45A mRNA was detected with specific probes for its 5’ and 3’ untranslated regions (5’UTR and 3’UTR). Positions of the 28S and 18S are indicated. Loading was controlled by detection of 28S rRNA. The relative expression of GADD45A mRNA was normalized to that of 28S rRNA. Then, relative GADD45A mRNA levels in conditions of DIS3L2 knockdown (KD) were normalized to those at control conditions (LUC KD), as indicated below each autoradiograph.

### The efficiency of DIS3L2-mediated decay increases in the presence of TUTases

To further proceed on understanding how DIS3L2 is involved in the degradation of NMD-targets, we took advantage of several studies showing that DIS3L2 preferentially acts on mRNA decay after an addition of untemplated uridine residues to the mRNA 3’-end [37–40,42]. Since mRNA uridylation is achieved mainly by the TUTases TUT4 and TUT7 [40], we performed experiments to access the involvement of these TUTases in the DIS3L2-mediated mRNA decay of NMD-targets. With this purpose, we performed KDs for TUT4, TUT7, TUT4+TUT7, DIS3L2+TUT4, DIS3L2+TUT7 and DIS3L2+TUT4+TUT7 (Fig 5A-C). Then, we accessed the mRNA levels of SMG5, SMG1, SLC1A3, GADD45A, and ATF3 mRNAs by RT-qPCR. Results were normalized to those obtained in conditions of LUC depletion and further compared to those obtained upon DIS3L2 knockdown (Fig 5D-H). Concerning the SMG5 mRNA (Fig 5D), none of the KDs showed a significant effect in the mRNA levels. This result is consistent with results obtained previously (Fig 2B) showing that SMG5 is not a target of DIS3L2. Regarding the SMG1 mRNA (Fig 5E), as expected, we observed a significant 2.7-fold increase in its mRNA levels upon DIS3L2 KD. Silencing TUTases 4 and 7 does not substantially affect its mRNA levels. Regarding the double KD of DIS3L2 with each of the TUTases (DIS3L2+TUT4 or DIS3L2+TUT7), we observed a 2.2- and 2.3-fold increase in the SMG1 mRNA levels, comparable to the levels obtained with DIS3L2 single KD. Interestingly, the DIS3L2+TUT4+TUT7 triple KD abolishes the accumulation of SMG1 mRNA levels (1.2-fold change comparing to LUC control conditions). This constitutes a significant 2.3-fold decrease in the SMG1 mRNA levels comparing to the cells with DIS3L2 single KD. Thus, the accumulation of SMG1 mRNA upon DIS3L2 KD seems to be dependent on the TUTases activity. Considering the SLC1A3 mRNA (Fig 5F), we observed a significant 2.4-fold increase in the SLC1A3 mRNA levels upon DIS3L2 KD. Silencing TUT4, TUT7 and both TUT4+TUT7 induces a 1.5-, 2.3-, and 1.9-fold increase in mRNA levels compared with those at control conditions. Considering the double KD of DIS3L2 together with TUT4 or TUT7, we observed a 2.1- and 1.4-fold increase in mRNA levels. Once more, the DIS3L2+TUT4+TUT7 triple KD abolishes the accumulation of SLC1A3 mRNA levels (1.1-fold change comparing to control conditions), which is a significant 2.2-fold decrease comparing to its levels in cells with the DIS3L2 single KD. Again, these results may indicate the need of TUTases activity for DIS3L2-mediated degradation. Considering the GADD45A mRNA (Fig 5G), we also observed a significant 5.2-fold increase in its mRNA levels upon DIS3L2 KD. Silencing TUT4, TUT7 or both TUT4+TUT7 does not substantially affect the GADD45A mRNA levels. Considering the double KD of DIS3L2 together with TUT4 or TUT7, we observed a 4.4- and 3.5-fold increase in the GADD45A mRNA levels, respectively. Interestingly, the DIS3L2+TUT4+TUT7 triple KD also abolishes the accumulation of GADD45A mRNA levels (1.2-fold change comparing to normal conditions), which constitutes a significant 4.3-fold decrease comparing to the levels observed in cells depleted of DIS3L2. Thus, the accumulation of GADD45A mRNA upon DIS3L2 depletion is also dependent on the TUTases activity. Regarding the ATF3 mRNA (Fig 5H), we observed a significant 4.7-fold increase in its mRNA levels upon DIS3L2 KD. Silencing TUT4, TUT7 or both, does not affect substantially the ATF3 mRNA levels. Considering the condition of double KD of DIS3L2 together with TUT4 or TUT7, we observed a 4.0- and 2.1-fold increase in the ATF3 mRNA levels. The DIS3L2+TUT4+TUT7 triple KD also abolishes the accumulation of ATF3 mRNA levels (1.2-fold change comparing to control conditions); this is a significant 3.9-fold decrease in mRNA levels comparing to those in cells with DIS3L2 single KD. Thus, also for this transcript, the accumulation of mRNA upon DIS3L2 KD is dependent on the TUTases activity. These results suggest that uridylation by TUT4/TUT7 marks SMG1, SLC1A3, GADD45A, and ATF3 mRNAs for degradation by DIS3L2.

**Fig 5.**
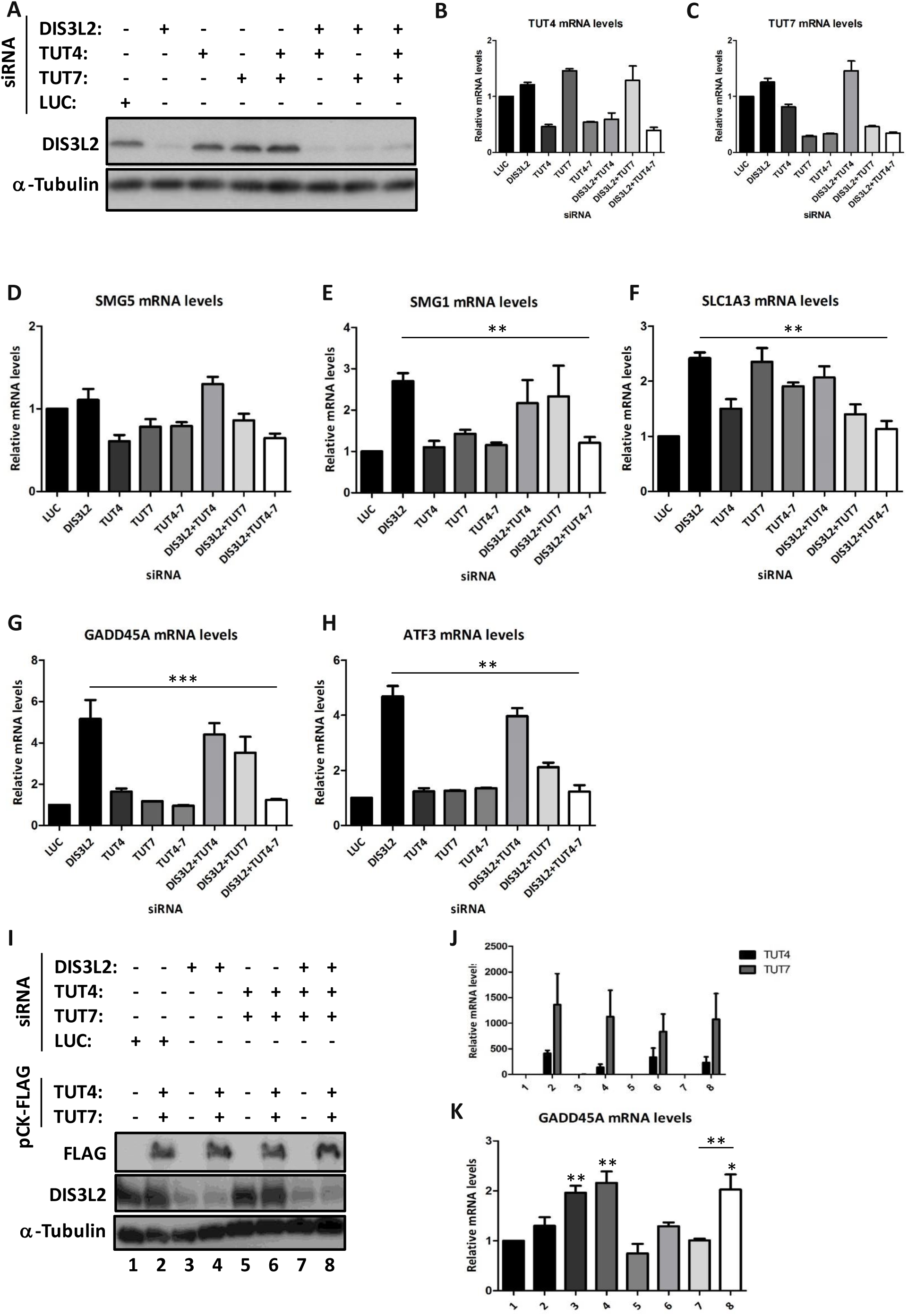
The efficiency of DIS3L2-mediated decay depends on uridylation by TUTases 4 and 7. (A) Representative Western blot analysis of HeLa cells extracts transfected with (+) or without (-) control Luciferase (LUC), DIS3L2, TUT4, TUT7, TUT4+TUT7, DIS3L2+TUT4, DIS3L2+TUT7 or DIS3L2+TUT4+TUT7 siRNAs. Protein levels present in the cell extracts were analyzed by Western blot for DIS3L2 and α-tubulin (loading control) to monitor DIS3L2 knockdown. Identification of each band is indicated to the left of the gel image. (B, C) RT-qPCR analysis of the TUT4 or TUT7 mRNA levels expressed in HeLa cells treated with control Luciferase (LUC), DIS3L2, TUT4, TUT7, TUT4+TUT7 (TUT4-7), DIS3L2+TUT4, DIS3L2+TUT7, or DIS3L2+TUT4+TUT7 (DIS3L2+TUT4-7) siRNAs, showing efficiency of TUT4 or TUT7 knockdown, respectively in (B) or (C). (D-H) mRNA levels of natural NMD-targets (SMG5, SMG1, SLC1A3, GADD45A, and ATF3) were determined by RT-qPCR using primers specific for each case (as indicated above each histogram), and for glyceraldehyde-3-phosphate dehydrogenase (GAPDH) gene. Histograms represent fold-change of each studied NMD-target in UPF1- or DIS3L2-depleted HeLa cells, relative to its level at Luciferase (LUC) siRNA-treated cells, arbitrarily set to 1. Average and standard deviation (SD) of three independent experiments corresponding to three independent transfections are shown in the histograms. Asterisks (*) indicate statistical significance relative to the control conditions for each NMD-target. *P<0.05, **P<0.01. (I) Representative Western blot analysis of HEK293 extracts transfected with (+) or without (-) siRNAs targeting LUC, DIS3L2, TUT4+TUT7, DIS3L2+TUT4+TUT7, with (+) or without (-) expression of pCK-FLAG-TUT4+pCK-FLAG-TUT7 vectors. Protein levels present in the cell extracts were analyzed by Western blot for FLAG (monitor TUTases expression), DIS3L2 (monitor DIS3L2 knockdown) and α-tubulin (loading control). Identification of each band is indicated to the left of the gel image. (J) RT-qPCR analysis of the TUT4 or TUT7 mRNA levels expressed in HEK293 cells treated with the same conditions as specified for Figure 5I, showing efficiency of TUT4 or TUT7 knockdown. (K) mRNA levels of the natural GADD45A NMD-target were determined by RT-qPCR using specific primers as above (legend to figure 5D-H). Unless otherwise stated, asterisks (*) indicate statistical significance relative to the mRNA levels in DIS3L2 knockdown condition. \**P*<0.05, \*\**P*<0.01, *** *P<*0.001.

To further confirm that TUT4 and TUT7 impact levels of DIS3L2/NMD-targets, we transfected HEK293 cells with plasmids expressing FLAG-TUT4 and FLAG-TUT7 (full-length human TUT4 and full-length human TUT7), simultaneously with siRNAs for LUC, DIS3L2, TUT4+TUT7 or DIS3L2+TUT4+TUT7 (Fig 5I-J). Then, we accessed the levels of GADD45A mRNA by RT-qPCR. Results were normalized to those obtained in conditions of LUC depletion (Fig 5K: line 1). Data show that overexpression of wild-type TUT4 and TUT7 does not significantly affect levels of GADD45A mRNA expressed in control conditions (lane 2 *versus* lane 1), or in conditions of DIS3L2 or TUT4/7 knockdown (lane 4 *versus* 3 and 6 *versus* 5, respectively). Nevertheless, under conditions of DIS3L2+TUT4+TUT7 knockdown, overexpression of TUT4 and TUT7 led to a significant increase in levels of GADD45A mRNA (Fig 5K: lane 8 *versus* 7), becoming comparable to those observed under DIS3L2 knockdown (Fig 5K: lane 8 *versus* 3). Taken together, our data show that DIS3L2 function in the degradation of NMD-targets depends on the activity of both TUT4 and TUT7.

To unequivocally show that uridylation is occurring at the 3’-end of the DIS3L2/NMD-targets, we analyzed the termini of the GADD45A mRNA 3’UTR by performing a 3’ rapid amplification of cDNA ends (3’-RACE) followed by Sanger sequencing of the GADD45A mRNAs expressed in cells transfected with LUC siRNA (control), DIS3L2, or DIS3L2+XRN1 siRNAs (Fig 6 and Fig S4). DIS3L2+XRN1 double depletion was also included in our analysis as XRN1 KD hampers 5’-3’ decay pathway, and thus mRNA decay might rely more on the 3’-5’ pathway and should be more sensitive to effects over DIS3L2 activity. We observed that in control conditions, GADD45A mRNAs are not uridylated (Fig 6 and Fig S4). In DIS3L2-depleted cells, we found a low but consistent frequency of uridylation (U_1_ to U_2_). In conditions of XRN1+DIS3L2 double KD, a much higher frequency of oligo(U) tails (U_3_ to U_15_) was detected.

**Fig 6.**
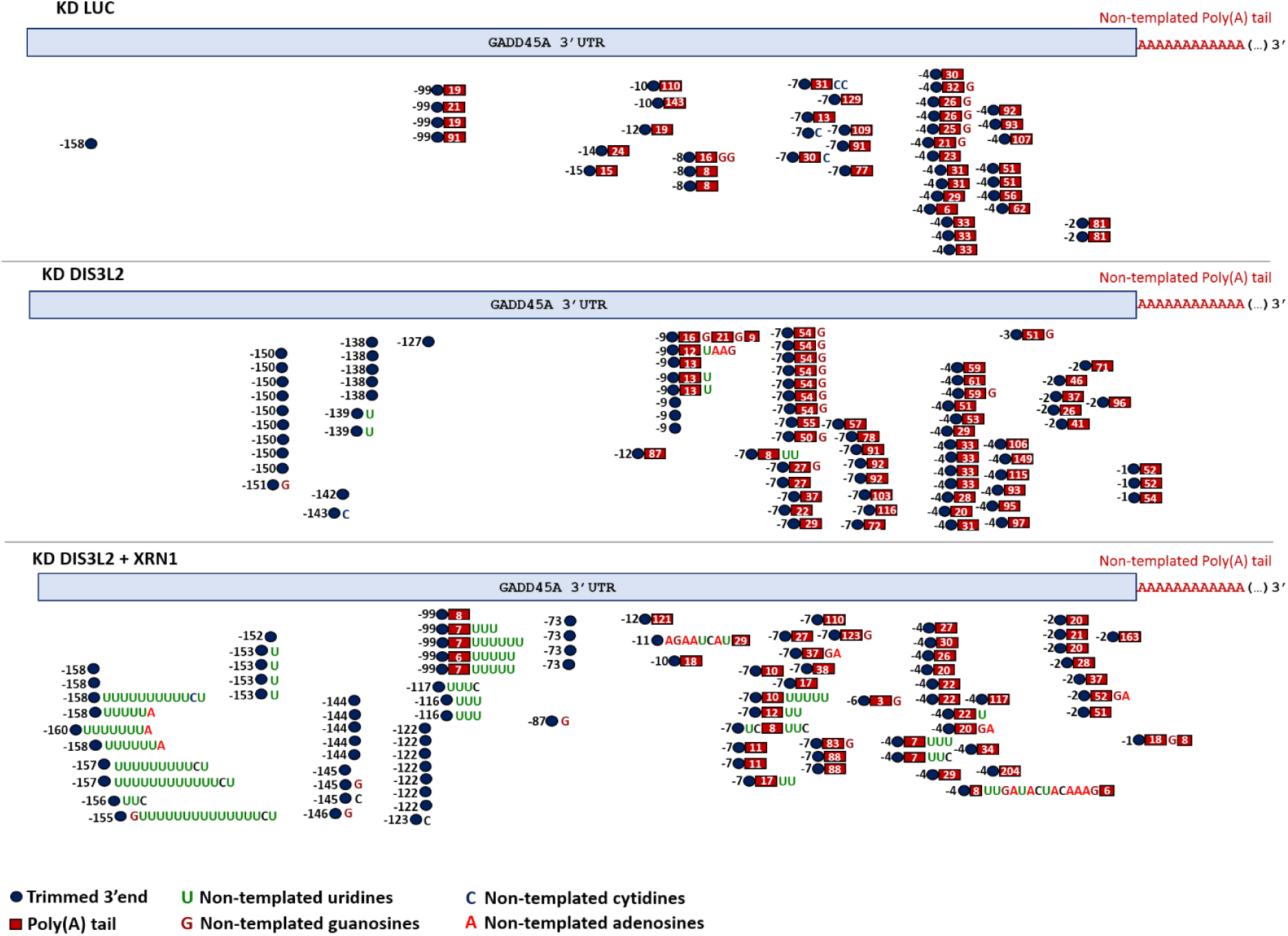
Depletion of XRN1+DIS3L2 in HeLa cells results in the accumulation of trimmed and uridylated GADD45A transcripts. Schematic representation of GADD45A mRNA uridylation frequency in LUC, DIS3L2 and DIS3L2+XRN1-depleted HeLa cells. Each blue circle represents the GADD45A mRNA 3’ end. The negative values upstream of the blue circle indicate the 3’ end position in the 3’UTR if the mRNA was trimmed, considering that the point zero is the polyadenylation site of GADD45A mRNA. Red squares indicate the length of the poly(A) tail added after trimming. Non-templated uridine residues detected at the 3’-end are indicated by green Us, the non-templated cytidine residues by blue Cs, the non-templated guanosine residues by dark red Gs, and non-templated adenosine residues by red As. The 3’ ends detected also revealed the existence of heterogeneous non-templated nucleotide additions.

A common feature of all the sequenced transcripts was the trimming of GADD45A mRNA within the 3’UTR. As we can see in Fig 6 and S4, we could find trimmed intermediaries, with and without a poly(A) tail, and in both cases, it was observed the addition of non-templated Us (except under the control conditions). Interestingly, the longest U-tails, found mostly on the XRN1+DIS3L2 double KD condition, were present in the non-adenylated and smallest 3’-trimmed transcripts. In the case of uridylated transcripts with a poly(A) tail, ranging between 0-22 nts, the U-tails are shorter. This goes in agreement with Chang *et al*. [54] that reported the existence of U-tails attached to short poly(A) tails (<25 nts).

According to data previously published [54, 55], we also observed terminal guanylation in the 3’-ends of transcripts, with a similar prevalence in all the conditions, and in both non-adenylated and adenylated transcripts. In agreement with the data reported by Chang *et al*. and Lim *et al*. [54,55] indicating that G-tails are found mainly on longer poly(A) tails (>40 nts), our adenylated transcripts with non-templated G-addition had an A-tail in the range of 3-83 nts. Citidylation was also found in all the conditions tested, but less frequently. Its occurrence seems to be independent of the poly(A) tail, as reported by Chang *et al*. [54]. Indeed, non-templated cytidine addition was also observed in some uridylated RNAs.

In the DIS3L2 and XRN1+DIS3L2 KD conditions, 3’-ends also revealed the existence of heterogeneous non-templated nucleotide additions. In the XRN1+DIS3L2 KD condition, were also found cases of A- or C-addition after U-tails. Together, our findings support the role of DIS3L2 and uridylation in the NMD pathway.

### DIS3L2 regulates the levels of a subset of UPF1-targets

Having showed that DIS3L2 regulates some specific NMD-targets, we decided to address if this role could represent a wider regulatory network. For this, we took advantage of publicly available data from transcriptome profiling of cells depleted of DIS3L2 or UPF1 [11,34]. In order to have a bona fide comparison between datasets, RNA sequencing approaches were chosen in the same cell type, HeLa cells, as well as the same silencing-method by siRNAs. Moreover, transcriptomic raw data upon UPF1 KD was retrieved and differentially expressed genes were detected using the same algorithm as used for the DIS3L2 KD by Lubas and colleagues [11] (Table S4). After cross-referencing the obtained gene sets, we found that 75 out of 681 (about 11%) upregulated genes after DIS3L2 KD are within the pool of upregulated transcripts detected in UPF1-depleted cells, supporting a functional link between DIS3L2 and NMD.

## DISCUSSION

In this work, our data support the involvement of DIS3L2 in NMD, in human cells. Indeed, we show that some NMD-targets are highly stabilized in DIS3L2-depleted HeLa cells. In accordance, our data indicate that DIS3L2 and the NMD key player, UPF1, function in the same pathway. Trying to dissect the mechanism through which DIS3L2 may be involved in NMD, we show that DIS3L2 activity is increased in the presence of TUTases 4/7. This goes in agreement with the unprecedented property of this enzyme of preferentially acting on uridylated RNAs [38]. Knowing that DIS3L2 does not interact with the exosome [34,38], our data indicate that NMD can involve a 3’-5’ cytoplasmic RNA degradation pathway independently from the exosome, through the DIS3L2 3’-5’ activity; thus, this new mRNA decay pathway can be added as a new branch of NMD. Reinforcing our results showing that DIS3L2 is involved in the degradation of specific natural NMD-targets, such as SMG1, SLC1A3 and GADD45A, Vanacova and co-workers [42] identified the SMG1 transcript as one uridylated mRNA targeted by DIS3L2. Furthermore, Kim and co-workers [40,54] showed that SLC1A3 mRNA is uridylated by TUT4/7. More recently, a study was also published showing that uridylation and DIS3L2 can mediate the degradation of NMD-targets [56]. Together, these data strongly support a prominent role for DIS3L2 in the degradation of NMD-targets. Interestingly, DIS3L2 has been associated with other mRNA decay pathways, as the general mRNA decay and the ARE-mediated decay [34,35,37]. Taking together, we can assume that DIS3L2 is not exclusive of a specific mRNA decay pathway, but it might be dependent on molecular interactions and other components as cofactors to target a wide variety of RNA substrates.

It has been shown that the NMD pathway is initiated by the central NMD factor UPF1, which recruits the endonuclease SMG6 and the deadenylation-promoting SMG5-SMG7 complex to the mRNA [28,29]. However, the extent to which SMG5-SMG7 and SMG6 contribute to the degradation of NMD-substrates is not completely understood. Lately, Ottens and colleagues (2017) have shown that endogenous transcripts can have NMD-eliciting features at various positions, including uORFs, PTCs, and long 3’UTRs [8]. These authors found that NMD-substrates with PTCs undergo constitutive SMG6-dependent endocleavage, rather than SMG7-dependent exonucleolytic decay, and in our study, this type of NMD-substrate behaves as DIS3L2-resistent. In contrast, the turnover of NMD-substrates containing uORFs and long 3’UTRs involves both SMG6- and SMG7-dependent endo- and exonucleolytic decay, respectively [8]. In our work, we observe that DIS3L2 functions in the decay of natural NMD-targets in a transcript-specific manner. Regarding the DIS3L2/NMD-targets here studied, the analysis of some of their features (including NMD-eliciting features; Table 1) indicates that DIS3L2-sensitive transcripts have common features with those that are DIS3L2-resistant (Table 1). In addition, motif search did not reveal a particular motif enriched in the 3’UTRs. Thus, what makes a natural NMD-target a DIS3L2-target is not yet clear. We can speculate about the function of regulatory elements and/or molecular interactions, as well as mRNA structure and context, but it remains to be identified the exact feature that makes a NMD-target a substrate for DIS3L2.

**TABLE 1.** Some features of the studied NMD-targets.

| **Endogenous NMD target** | **DIS3L2 target** | **3'UTR length (nts)** | **uORF length\* (codons)** | **3'UTR intron** | **3'UTR GC content (%)** |
| --- | --- | --- | --- | --- | --- |
| SMG5 | No | 1377 | 31 | No | 59 |
| ARFRP1 | No | 1800 | No | No | 64 |
| BAG1 | No | 2745 | 17; 275; 231 | No | 45 |
| ANTXR1 | No | 3840 | No | No | 40 |
| GABARAPL1 | No | 1291 | No | No | 45 |
| SLC7A11 | No | 7800 | 20 | No | 33 |
| SMG1 | Yes | 4716 | 7 | No | 48 |
| SLC1A3 | Yes | 2063 | No | No | 39 |
| GADD45A | Yes | 582 | No | No | 31 |
| GADD45B | Yes | 682 | No | No | 55 |
| PLXNA1 | Yes | 3375 | No | No | 60 |
| ATF3 | Yes | 1221 | 96 | No | 47 |

nts: nucleotides; uORF: upstream open reading frame; \*for uORFs initiated by an uAUG

Lim and colleagues have shown that two TUTases, TUT4 and TUT7, uridylate mRNAs with short A-tails at the 3’-end [40]. Moreover, it was shown that DIS3L2 preferentially binds to U-tails and this is involved in the turnover of some miRNAs and mRNAs in a uridylation-dependent manner [34,38,39,42,57]. More recently, the connections between DIS3L2 and uridylation in RNA degradation were expanded with the involvement of TUTases and DIS3L2 in template-dependent miRNA degradation (TDMD) and mRNA degradation in apoptotic cells [41,58]. In addition, the determination of DIS3L2 structure revealed that substrate recognition involves three uracil-specific interactions spanning the first 12 nucleotides of an oligoU-tailed RNA. This explained how DIS3L2 recognizes, binds and processes preferentially oligoU-tailed RNA [57]. Considering the results shown in Fig 5, we can assume that the DIS3L2-mediated decay of SMG1, GADD45A, SLC1A3 and ATF3 mRNAs depends on the activity of both TUT4 and TUT7. This suggests that uridylation is also a mark for degradation of NMD-targets that subsequently are degraded by DIS3L2 from 3’-to-5’. Thus, we propose a model in which 3’-end uridylation marks transcripts for DIS3L2 degradation. When DIS3L2 is absent these transcripts accumulate, however, when uridylation is abrogated, or at least inhibited, these transcripts may become vulnerable to other exoribonucleases being expressed at levels comparable with those observed at normal conditions. Besides this, a previous work from Thomas and colleagues (2015), in which they show that some of the uridylated mRNAs by TUT4 and TUT7 are not decapped, suggest a predominance of 3’-to-5’ decay upon uridylation [41]. In addition, our results also show that DIS3L2/NMD-targets do not need to be cleaved by SMG6 (Fig 4 and S3), supporting the idea that DIS3L2 uridylated targets are full-length. Moreover, our results demonstrate redundancy between TUT4 and TUT7, as DIS3L2+TUT4 or DIS3L2+TUT7 double depletions do not abolish the accumulation of the DIS3L2-targeted mRNAs. We hypothesize that both TUTases may function as “fail-safe proteins”.

In this work, the prevalence of uridylation and the position where it occurs inside GADD45A mRNA was confirmed by 3’-RACE analysis (Fig 6 and S4). In fact, we observed that at control conditions (Luciferase siRNA-treated cells), GADD45A mRNAs are not uridylated. In DIS3L2 depleted cells, we observed mono- and di-uridylation in non-adenylated and oligoadenylated 3’-end trimmed transcripts. Of note, in the XRN1+DIS3L2 double KD, the oligo-uridylation (U_3_ to U_15_) was the most frequent 3’-end modification observed, being the majority of the transcripts 3’-end trimmed molecules without poly(A) tail. These results suggest that uridylation marks these NMD-target intermediates for degradation by DIS3L2. When DIS3L2 is not able to degrade these transcripts, the TUTases keep uridylating the mRNA, until achieve oligo(U) tails of at least 15 Us. These oligo(U) tails might activate the decapping, allowing the 5’-to-3’ decay by XRN1, as previously observed in yeast [59]. Thus, the uridylation-dependent mRNA decay might function as a fail-safe mechanism to stimulate decapping and further 5’-to-3’ decay by XRN1. Our results also suggest that oligouridylated NMD-targets are subject to degradation by multiple factors, as previously described for deadenylated mRNAs [40,54]. Furthermore, according with previous works from Narry Kim group [54,55], we also observed non-templated guanylation and citidylation. Conversely to uridylation, these non-templated additions were reported to increase mRNA stability [55]. In addition and according to our data, it was recently shown that phosphorylated UPF1, which is the active form of this essential NMD factor, binds predominantly deadenylated mRNA decay intermediates, instead of SMG6 endonucleolytic cleavage intermediates. It activates NMD cooperatively from 5′ and 3′ ends, being some of the 3’-ends subject to the addition of non-templated uridines by TUT4 and TUT7 and degraded by DIS3L2 [56].

In conclusion, our work shows that DIS3L2 is an important novel factor in NMD, degrading specific natural NMD-targets independently of SMG6 and the exosome. Therefore, this 3′-to-5′ degradation pathway should be considered as an alternative in NMD. Future investigation will surely uncover the details regarding the mechanism, and specific transcripts’ features, involved in triggering NMD-substrates for DIS3L2 degradation.

## ACKNOWLEDGEMENTS

This work was partially supported by Fundação para a Ciência e a Tecnologia (FCT), Portugal (PTFC/BIM-MEC/3749/2014 to LR and UID/MULTI/04046/2013 centre grant to BioISI). PJdC and JFG-M are recipients of a fellowship from BioSys PhD programme (SFRH/BD/52495/2014 to PJdC, SFRH/BD/52492/2014 to HAS, and PD/BD/142898/2018 to JFG-M) and JM is a posdoc fellow (SFRH/BPD/98360/2013) from FCT.

Work at ITQB NOVA was financially supported by: Project LISBOA-01-0145-FEDER-007660 (Microbiologia Molecular, Estrutural e Celular) funded by the European Regional Development Fund (FEDER) through COMPETE2020 - Programa Operacional Competitividade e Internacionalização (POCI) and by national funds through FCT: project PTDC/BIA-MIC/1399/2014 to CMA and project PTFC/BIM-MEC/3749/2014 to SCV. SCV was financed by program IF of FCT [ref. IF/00217/2015]. MS was financed by an FCT grant [SFRH/BPD/109464/2015]. We thank Dr. V. Narry Kim from Seoul National University (Seoul, Korea), who kindly provided us with the pCK-FLAG-TUT4 and pCK-FLAG-TUT7 vectors.

