## Supplemental data for "A role for DIS3L2 over human nonsense-mediated mRNA decay targets"

### Supplementary Information

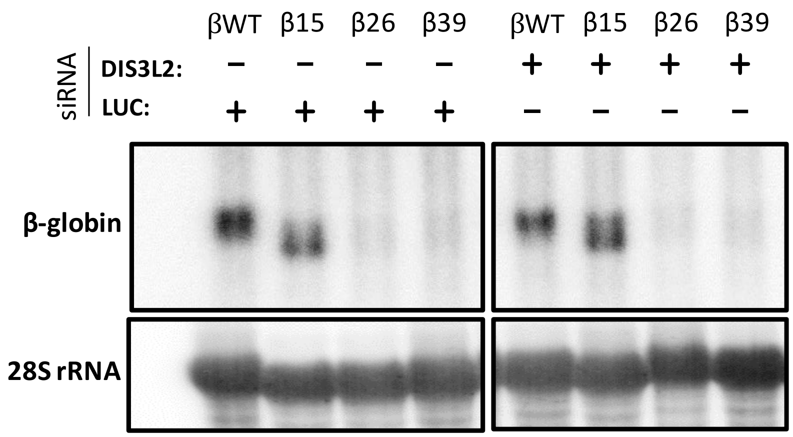


**Figure S1. DIS3L2 does not affect human β-globin transcripts.** Northern blot analysis of total RNA obtained from HeLa cells treated with (+) or without (-) siRNAs targeting DIS3L2. Twenty micrograms of total RNA were separated under denaturing conditions on an agarose MOPS/formaldehyde gel (1.5%) and human β-globin variants: βWT, β15, β26 and β39 were detected with a specific probe for exon 1. Loading was controlled by detection of 28S rRNA.

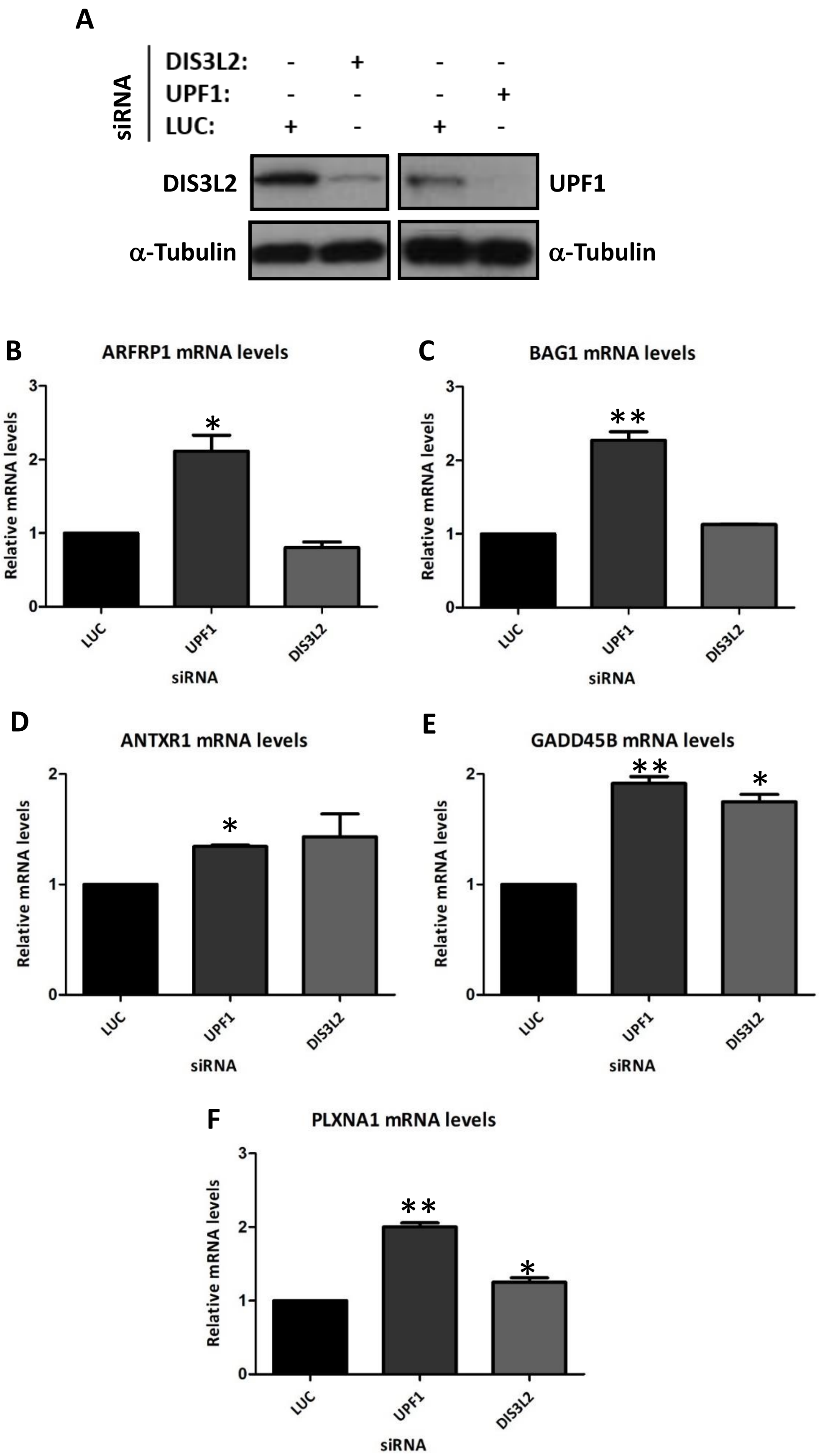
A
siRNA
DIS3L2: - + - -
UPF1: - - - +
LUC: + - + -
DIS3L2
UPF1
α-Tubulin
α-Tubulin
B
ARFRP1 mRNA levels
Relative mRNA levels
LUC
UPF1
DIS3L2
siRNA
C
BAG1 mRNA levels
Relative mRNA levels
LUC
UPF1
DIS3L2
siRNA
D
ANTXR1 mRNA levels
Relative mRNA levels
LUC
UPF1
DIS3L2
siRNA
E
GADD45B mRNA levels
Relative mRNA levels
LUC
UPF1
DIS3L2
siRNA
F
PLXNA1 mRNA levels
Relative mRNA levels
LUC
UPF1
DIS3L2
siRNA

**Figure S2. UPF1 and DIS3L2 can act in common transcripts.** (A) Representative Western blot analysis of HeLa cells extracts transfected with (+) control Luciferase siRNA (LUC siRNA), or with siRNAs targeting the human UPF1 or DIS3L2. Protein levels present in the cell extracts were analyzed by Western blot for UPF1, DIS3L2 and α-tubulin (loading control) to monitor knockdown of UPF1 and DIS3L2, respectively. Identification of each band is indicated to the left and right of the gel image. (B-F) mRNA levels of natural NMD-targets (ARFRP1, BAG1, ANTXR1, GADD45B and PLXNA1) were determined by RT-qPCR using primers specific for each case (as indicated above each histogram), and for glyceraldehyde-3-phosphate dehydrogenase (GAPDH) gene. Histograms represent fold-change of each studied NMD-target in UPF1- or DIS3L2-depleted HeLa cells, relative to its level at Luciferase (LUC) siRNA-treated cells, arbitrarily set to 1. Average and standard deviation (SD) of three independent experiments corresponding to three independent transfections are shown in the histograms. Asterisks (*) indicate statistical significance relative to the control conditions for each NMD-target. *$P<0.05$, **$P<0.01$.

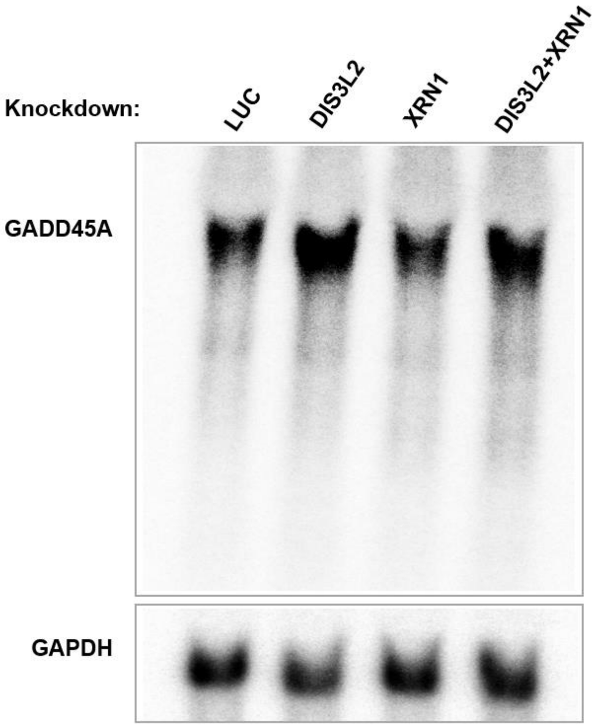


**Figure S3. GADD45A decay intermediates were not detected.** Northern blot analysis of total RNA obtained from HeLa cells treated with (+) or without (-) siRNAs targeting DIS3L2, XRN1, or DIS3L2+XRN1. Total RNA was separated under denaturing conditions on an agarose MOPS/formaldehyde gel (1.2%) and GADD45A mRNA was detected with a specific probe for its 3'UTR region (Lykke-Andersen et al. 2014. Genes Dev. 28: 2498-517. doi: 10.1101/gad.246538). Position of the full-length GADD45A mRNA is indicated. Loading was controlled by detection of GAPDH mRNA.

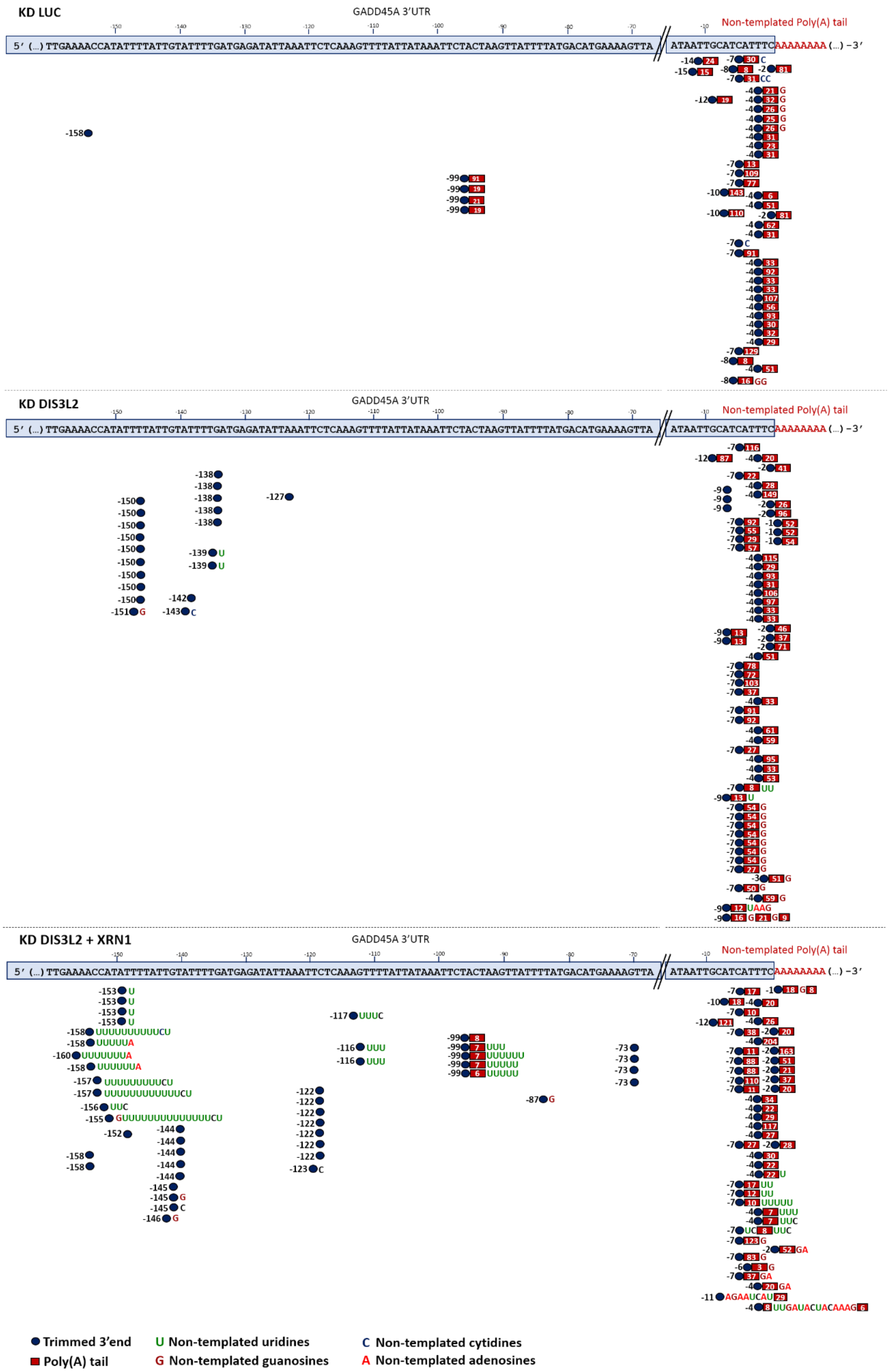
KD LUC
GADD45A 3'UTR
Non-templated Poly(A) tail
5′ (...) TTGAAAACCATATTTTATTGTATTTTGATGAGATATTAAATTCTCAAAGTTTTATTATATAAATTCTACTAAGTTATTTTATGACATGAAAAGTTA
ATAATTGCATCATTTCAAAAAAAA (...) −3′
KD DIS3L2
GADD45A 3'UTR
Non-templated Poly(A) tail
KD DIS3L2 + XRN1
GADD45A 3'UTR
Non-templated Poly(A) tail
Trimmed 3'end
Poly(A) tail
U Non-templated uridines
G Non-templated guanosines
C Non-templated cytidines
A Non-templated adenosines

**Figure S4. Depletion of XRN1+DIS3L2 in HeLa cells results in the accumulation of trimmed and uridylated GADD45A transcripts.** Representation at scale of uridylation frequency of GADD45A mRNA expressed in LUC, DIS3L2 and DIS3L2+XRN1-depleted HeLa cells. Each blue circle represents the GADD45A mRNA 3' end. The negative values upstream of the blue circle indicates the exact 3' end position in the 3' UTR if the mRNA was trimmed, considering that the point zero is the polyadenylation site of GADD45A mRNA. Non-templated uridine residues detected at the 3'-end are indicated by green Us, the non-templated cytidine residues by blue Cs, the non-templated guanosine residues by dark red Gs, and non-templated adenosine residues by red As. The 3' ends detected also revealed the existence of heterogeneous non-templated nucleotide additions.

### Supplementary Materials and Methods

#### Northern blot analysis

For Northern blot analysis, equal amounts of human total RNA samples were separated under denaturating conditions in an agarose MOPS/formaldehyde gel following the procedure described (Malecki et al. 2013. EMBO J. 32: 1842-54. doi: 10.1038/emboj.2013.63). Briefly, the RNA was transferred onto Hybond-N+ membranes by capillarity using 20×SSC as transfer buffer, and UV cross-linked to the membrane immediately after. Membranes were then hybridized, in PerfectHyb Buffer (Sigma) for 16 hours, with the respective riboprobe at 68°C or oligoprobe at 43°C. After hybridization, membranes were rinsed at RT in a 2X SSC/0.1% SDS solution, followed by washing in three subsequent 15 minutes steps in SSC (2X, 1X or 0.5X, respectively)/0.1% SDS solutions at the hybridization temperature. Signals were visualized by Phosphor Imaging (FUJI TLA-5100 Series, Fuji) and analyzed using the ImageQuant software (GE Healthcare).

In Figure S1, the human β-globin riboprobe was specific for exon 1 region and 28S rRNA was detected using an oligoprobe. In Figure S2, GADD45A mRNA was detected with a riboprobe specific for 3'UTR region as described in (Lykke-Andersen et al. 2014. Genes Dev. 28: 2498-517. doi: 10.1101/gad.246538) and GAPDH was detected using an oligoprobe (see Supplementary Table 3).

**Supplementary Table 1.** Sequences of siRNAs used in this work.

| **Targeted gene** | **Sequence (5'→ 3')** |
|---|---|
| Luciferase (LUC) | CGUACGCGGAAUACUUCGA |
| DIS3L2 | GCACCAAACUUAGCUACGA |
| XRN | GGGAUCUGGAAAGAUGCAAUACUUU |
| TUT4 (ZCCHC11) | GGAUUUGGAUUUCGUGAUA |
| siTUT4_1 | GGUUGCUUCAGACUUUAUA |
| TUT7 (ZCCHC6) | GGCUGGAAAUUAAACGUAU |
| siTUT7_4 | GAACAUGAGUACCUAUUUA |

**Supplementary Table 2.** Sequence of primers used for RT-qPCR analysis.

| **Gene** | **Orientation** | **Sequence (5'→3')** |
|---|---|---|
| HBB | Forward | GTGGATCCTGAGAACTTCAGGC |
| HBB | Reverse | CAGCACACAGACCAGCACGT |
| GAPDH | Forward | CCATGAGAAGTATGACAACAGCC |
| GAPDH | Reverse | GGGTGCTAAGCAGTTGGTG |
| SMG5 | Forward | CCCCTCATAGGATGCAAGAA |
| SMG5 | Reverse | ATCTGTGCCCAATCCATCTC |
| SLC7A11 | Forward | GGGCATGTCTCTGACCATCT |
| SLC7A11 | Reverse | TCCCAATTCAGCATAAGACAAA |
| GADD45A | Forward | GGAGGAATTCTCGGCTGGAG |
| GADD45A | Reverse | CGTTATCGGGGTCGACGTT |
| GABARAPL1 | Forward | GGCCAGTTCTACTTCTTAATCCGG |
| GABARAPL1 | Reverse | AGGTGCTCCCATCTGCTGGG |
| SMG1 | Forward | TGTGAGCAGGTTTTACACATTATGC |
| SMG1 | Reverse | CCAGAGGGTCGTACACAAAGG |
| GADD45B | Forward | ACAGTGGGGGTGTACGAGTC |
| GADD45B | Reverse | GGATGAGCGTGAAGTGGATT |
| SLC1A3 | Forward | TTCTCCTTTCCTGGGGAACT |
| SLC1A3 | Reverse | CCATCTTCCCTGATGCCTTA |
| TUT4 | Forward | AAAAGGGACCCAGTTTACTGTTG |
| TUT4 | Reverse | GTCCGATACGTCTTCAATTCCTG |
| TUT7 | Forward | ATAACACCAGGGAACTATGGGA |
| TUT7 | Reverse | CATTCATCCAAGCGGGTTGAC |
| DIS3L2 | Forward | ACCGCGAGAGCAACAAGCT |
| DIS3L2 | Reverse | GATCTTGTGGGCCACTGC |
| DIS3L1 | Forward | CTTTTGTTCGACTTCAAGGAGCC |
| DIS3L1 | Reverse | CCGTGGATTAGGATGTCACTGA |
| ATF3 | Foward | ATCACAAAAGCCGAGGTAGC |
| ATF3 | Reverse | TCTTGTTTCGGCACTTTGC |
| ARFRP1 | Forward | CTAAACATCGGCACTGTGGA |
| ARFRP1 | Reverse | ACTTGTCCCACAAAGACTGC |
| BAG1 | Forward | TGGGAAAAAGAACAGTCCACA |
| BAG1 | Reverse | TCCAGCTGGTCAGCTATCTTC |
| ANTXR1 | Forward | ATGCCTTGTGGGTCCTACTG |
| ANTXR1 | Reverse | GAGGTGTGGTAGGCGTTGTT |
| PLXNA1 | Forward | GTGAAGAACCACGACCACCT |
| PLXNA1 | Reverse | CGTGCTGAAGATGGTCTCAA |

**Supplementary Table 3.** Sequence of primers used for Northern blot and RACE analysis.

| Name | | Sequence |
|---|---|---|
| **Probe** | ***Primers for Northern blot probes*** | |
| | 28S rRNA Oligo | AACGATCAGAGTAGTGGTATTTCACC |
| GADD45A 5’UTR probe | GADD45A_RT_FW | GGAGGAATTCTCGGCTGGAG |
| | GADD45A 5’UTR T7 Promoter | GTTTTTTTTAATACGACTCACTATAGGCGTTATCGGGGTCGACGTT |
| GADD45A 3’UTR probe | GADD45A_inner_long_FW | GGAAAGTCGCTACATGGATCA |
| | GADD45A 3’UTR T7 Promoter | GTTTTTTTTAATACGACTCACTATAGGTGATTAACCCTTTCGGCTTTTC |
| β-globin probe | T7 Exon 1 Rv | GTTTTTTTTAATACGACTCACTATAGGCAGTAACGGCAGACTTCTCCTCAG |
| | 5’UTR Tex | ACATTTGCTTCTGACACAAC |
| ***Primers for RACE analysis*** | | |
| Linker Oligo | | 5’P-GATACTACCTCTATGAATTCTTTGCTAGCTACCTGAACTTATCAGA CCTAC-3’-$NH_3$ |
| BRevOligo | | GTAGGTCTGATAAGTTCAGGTAGCT |
| ARevOligo | | AGCAAAGAATTCATAGAGGTAGTATC |
| GADD45A External | | GTGTGCAGCCATAGTTTG |
| GADD45A Internal | | GCCGAAAGGGTTAATCATATTTG |
